# Cyanoexosortase B is essential for motility, biofilm formation and scytonemin production in a filamentous cyanobacterium

**DOI:** 10.1101/2024.12.02.626408

**Authors:** Gabriel A. Parrett, Daniel H. Haft, Maida Ruiz, Ferran Garcia-Pichel, Christopher C. Ebmeier, Douglas D. Risser

## Abstract

Exosortases are involved in trafficking proteins containing PEP-CTERM domains to the exterior of gram-negative bacterial cells. The role of these proteins in cyanobacteria, where such homologs are common, has not been defined. The filamentous cyanobacterium *Nostoc punctiforme* contains a single putative exosortase, designated cyanoexosortase B (CrtB), implicated by previous work in both the motility, and in the production of the UV-absorbing pigment, scytonemin. To determine the role of *crtB* in *N. punctiforme*, a *crtB*-deletion strain (Δ*crtB*) was generated. Δ*crtB* presented the loss of motility, biofilm formation, and scytonemin production. In the case of motility, the Δ*crtB* mutant exhibited a specific defect in the ability of hormogonia (specialized motile filaments) to adhere to hormogonium polysaccharide (HPS) and several PEP-CTERM proteins expressed in motile hormogonia were differentially abundant in the exoproteome of the wild type compared to the Δ*crtB* strain. These results are consistent with the hypothetical role of CrtB in the processing and export of PEP-CTERM proteins that play a critical role in stabilizing the interaction between the filament surface and HPS to facilitate motility and biofilm formation. In the case of scytonemin, the late biosynthetic steps of which occur in the periplasm and whose operon contains several putative PEP-CTERM proteins, Δ*crtB* failed to produce it. Given the abundance of putative PEP-CTERM proteins encoded in the *N. punctiforme* genome, and the fact that this study only associates a fraction of them with biological functions, it seems likely that CrtB may play an important role in other biological processes in cyanobacteria.

**Importance:** In gram-negative bacteria, exosortases facilitate the trafficking of proteins to the exterior of the cell where they have been implicated in stabilizing the association of extracellular polymeric substances (EPS) with the cell surface to facilitate biofilm formation and flocculation, but the role of exosortases in cyanobacteria has not been explored. Here, we characterize the role of cyanoexosortase B (CrtB) in the filamentous cyanobacterium *Nostoc punctiforme*, demonstrating that *crtB* is essential for motility, biofilm formation, and the production of the sunscreen pigment scytonemin. These findings have important implications for understanding motility and biofilm formation in filamentous cyanobacteria as well as efforts toward heterologous production of scytonemin in non-native hosts.

## Introduction

Filamentous cyanobacteria frequently exhibit gliding motility on solid surfaces (1). The benefits of this behavior include dispersal to new environments and migration in response to light signals, a phenomenon referred to as phototaxis (2). This allows the bacteria to colonize areas with sufficient light to support photosynthesis while avoiding high light environments that result in photodamage. Motility is also critical for the establishment of nitrogen-fixing symbioses with plants (3, 4). Additionally, motility in filamentous cyanobacteria is associated with biofilm formation (5–7) and facilitates the assembly of macroscopic aggregates (3, 8, 9). These aggregates are essentially self-assembled biofilms where the filaments collect and form a surface for accumulation of more filaments.

Motility in filamentous cyanobacteria is powered by rings of type IV pilus systems positioned at either end of the cell, adjacent to the septa that separate cells in the filament and requires the deposition of a motility-associated polysaccharide (10–12). The polysaccharide is thought to interface with the substratum and provide an attachment point for the T4P. Thus, the gliding motility of filamentous cyanobacteria relies on the production of a tube of polysaccharide through which the filament then pulls itself. Current understanding of the genetic underpinnings of the motility-associated polysaccharide production comes from studies on the genetically tractable model filamentous cyanobacterium *Nostoc punctiforme* ATCC29133 (PCC73102). Because, like most other heterocystous cyanobacteria, *N. punctiforme* only exhibits motility in specialized filaments termed hormogonia, its motility-associated polysaccharide was designated hormogonium polysaccharide (HPS). The precise chemical composition of HPS has not been resolved, but lectin-based methods indicate it contains fucose and galactose (10–12). Genetic analysis has identified several loci dispersed throughout the genome that encode genes involved in HPS production (11, 12). Most of the genes characterized code for glycosyl transferases presumed to be involved in assembling the sugar polymers. Genomic co-occurrence analysis indicates that this gene set is highly conserved in nearly all filamentous cyanobacteria (12), implying that most motile filamentous cyanobacteria employ an HPS-like polysaccharide for movement. The HPS export mechanisms, however, are yet to be fully determined. Some studies have suggested that it may be transported out of the cell via the type IV pilus systems (10). However, the presence of a genetic locus with genes encoding a Wzx/Wzy-type polysaccharide synthesis and export system is highly correlated to the presence of other HPS genes within cyanobacterial genomes (12). This type of correlational evidence implies that this system may be responsible for HPS secretion.

Intriguingly, this genetic locus also contains the cyanoexosortase B (*crtB*) gene (Npun_F0456), a gene that shares patterns of co-occurrence with other HPS genes, suggesting that it may also play a role in production/excretion of HPS (12). CrtB is a member of the exosortase family, a class of integral membrane proteins found in gram-negative bacteria that is typically involved in the processing of proteins that are either attached to the cell surface or released from the cell (13, 14). Target proteins for processing by exosortases contain an N-terminal signal sequence for transport across the cytoplasmic membrane via the general secretory pathway, and a PEP-CTERM domain at the C-terminus, consisting of Proline-Glutamate-Proline followed by a transmembrane alpha-helix and then a region enriched in basic amino acids (13, 14). The PEP-CTERM domain is thought to comprise the signal for processing by the exosortase. Aside from the N- and C-terminus, PEP-CTERM proteins are highly divergent among themselves, usually lack homology to proteins of known function, and are enriched in amino acids associated with glycosylation (13, 14). In most organisms, exosortases are found at genetic loci encoding genes involved in the synthesis and export of extracellular polysaccharides (EPS) (13, 14). PEP-CTERM proteins have been shown to be critical for stabilizing the interaction between EPS and the cell surface to facilitate floc formation in *Zoogloea resiniphila* and *Aquincola tertiaricarbonis* (16, 17). In *Flavobacterium johnsoniae*, the variant-type exosortase XrtF and its adjacent partner protein have been shown to be essential for biofilm formation and surface colonization, although target proteins for XrtF are not yet known (15). In cyanobacteria the *scy* locus, which is involved in production of the natural sunscreen pigment scytonemin, targeted for accumulation on the extracellular polysaccharide, encodes three putative PEP-CTERM proteins, ScyD, ScyE, and ScyF (18), with ScyE shown to be essential for scytonemin production in *N. punctiforme* (19).

The genomic association of *crtB* with genes thought to be involved in HPS production indicates that *crtB* and PEP-CTERM proteins may play a key role in the motility of filamentous cyanobacteria, while the presence of PEP-CTERM proteins encoded at the *scy* locus implicates a role for CrtB in scytonemin production. Here, using a combination of genetic and proteomic approaches in *N. punctiforme*, we demonstrate that *crtB* is essential for motility, biofilm formation and scytonemin production, and that it more generally influences the accumulation of extracellular PEP-CTERM proteins.

## Materials and Methods

### Strains and culture conditions

For a detailed description of the strains used in this study refer to Table S1. *N. punctiforme* ATCC 29133 and its derivatives were cultured in Allen and Arnon medium diluted four-fold (AA/4), without supplementation of fixed nitrogen, as previously described (20), with the exception that 4 and 10 mM sucralose was added to liquid and solid medium, respectively, to inhibit hormogonium formation (21). For hormogonium induction for phenotypic analysis, the equivalent of 30 μg chlorophyll *a* (Chl *a*) of cell material from cultures at a Chl *a* concentration of 10-20 μg ml^-1^ was harvested at 2,000 *g* for 3 min, washed two times with AA/4 and resuspended in 2 ml of fresh AA/4 without sucralose. For selective growth, the medium was supplemented with 50 μg ml^-1^ neomycin. *Escherichia coli* cultures were grown in lysogeny broth (LB) for liquid cultures or LB supplemented with 1.5% (w/v) agar for plates. Selective growth medium was supplemented with 50 μg ml^-1^ kanamycin, 50 μg ml^-1^ ampicillin, and 15 μg ml^-1^ chloramphenicol.

### Plasmid and strain construction

For a detailed description of the plasmids, strains, and oligonucleotides used in this study refer to Table S1. All constructs were sequenced to insure fidelity.

To construct plasmid pDDR539 for in-frame deletion of *crtB*, approximately 900 bp of flanking DNA on either side of the gene and several codons at the beginning and end of the gene were amplified via overlap extension PCR using primers NpF0456-5’-BamHI-F, NpF0456-5-OEP’-R, NpF0456-3’-OEP-F and NpF0456-3’-SacI-R, and cloned into pRL278 (22) as a BamHI-SacI fragment using restriction sites introduced on the primers.

To construct plasmid pGAP100, a mobilizable shuttle vectors containing *crtB* expressed from the *petE* promoter, the coding-region of *crtB* was amplified via PCR using primers NpF0456-BamHI-F and NpF0456-SacI-R and subsequently cloned into pDDR155 (3) as a BamHI-SacI fragment, replacing the *hmpA-gfp* coding region, using restriction sites introduced on the primers.

Gene deletion was performed as previously described (11) with *N. punctiforme* cultures supplemented with 4 mM sucralose to inhibit hormogonium development and enhance conjugation efficiency (23) (21). To construct UOP217, plasmid pDDR539 was introduced into wild type *N. punctiforme* ATCC29133.

### Motility assays

Plate and time lapse motility assays were performed as previously described (10). Briefly, for plate motility assays, colonies were transferred from AA/4 solid medium (1% noble agar) containing 5% sucrose, to suppress hormogonium development, to the surface of AA/4 solid medium (0.5% noble agar) without sucrose. Plates were incubated for 48 h under light. For time lapse motility assays, following standard hormogonium induction from liquid cultures, 2 ul of culture was spotted onto the surface of AA/4 solid medium (0.5% noble agar), overlayed with a cover slip, and imaged at 15s intervals. Both plate and time lapse motility assays were images with a Leica SD9 dissecting microscope equipped with a Leica Flexcam C3 camera controlled by Leica LAS X software. All assays were repeated in triplicate, with representative images and videos depicted.

### Immunoblot and lectin blot analysis

Preparation of *N. punctiforme* cell material, protein extraction and detection of PilA, RbcL, and HmpD, by immunoblot analysis was performed as previously described (16). Briefly, total cellular protein was extracted from cell material equivalent to 30 μg Chl*a* following standard protocols (16), lysate containing extracted proteins was separated on a 4-12% SDS-PAGE gel, and then transferred to a nitrocellulose membrane. Polyclonal antibodies raised against PilA (4), HmpD (32), and RbcL (33) were used at a 1:10,000 dilution, followed by a 1:20,000 dilution of an HRP-conjugated anti-rabbit secondary antibody (Chemicon). Lectin blot analysis to detect soluble HPS was performed as previously described (10). Briefly, 100 μl of cell-free culture medium was vacuum transferred to a nitrocellulose membrane and HPS was detected using biotinylated Ulex Europaeus Agglutinin I (UEA) (Vector Laboratories) following standard protocols (10).

### Immunofluorescence and fluorescent lectin staining

Detection of surface PilA and cell-associated HPS by immunofluorescence and fluorescent lectin staining was performed as previously described (10) (16). Briefly, cells were fixed in 4% paraformaldehyde, followed by methanol and acetone fixation, and subsequently polyclonal α-PilA antibodies (4) and UEA-fluorescein (Vector Laboratories) were used to detect PilA and HPS respectively following standard protocols (4, 16).

### HPS complementation assays

To collect conditioned culture medium for HPS complementation assays, standard hormogonium inductions (as described above) were performed for the appropriate strains. 24 h following induction, the cultures were harvested by centrifugation at 2,000 x g for 3 min and the supernatant was transferred to a new tube. The supernatant was subsequently centrifuged at 16,000 x g for 10 min to ensure complete removal of cell material, and following centrifugation the supernatant was collected in a new tube and stored at 4°C. For HPS complementation assays, standard hormogonium inductions (as described above) were performed for the appropriate strains, only using 15 μg Chl *a* of cell material instead of 30 μg Chl *a*. Following the second wash with AA/4 medium, the cell material was resuspended in 1 ml of conditioned medium from the appropriate strain and incubated under light for 24 h. Subsequently, time-lapse microscopy was performed (as described above) to visualize motility of individual filaments.

### Biofilm assays

To perform biofilm assays, 2 mL of culture containing 30 μg Chl *a* of cell material was transferred to a well of a 24 well plate and incubated under light with shaking at 120 rpm for 5 days. Daily, the cultures were vigorously pipetted to disperse the large colonial aggregate that forms in cultures producing hormogonia. On the fifth day, the cultures were removed from the wells, the wells were washed twice with 2 mL of AA/4 medium and subsequently images were taken. After imaging, the wells were washed with 1 mL of methanol to extract the Chl *a* from cell material adhering to the well. Quantification of Chl *a* was subsequently performed by measuring absorbance at OD_665_.

### Exoproteome analysis

Culture conditioned medium for exoproteome analysis was prepared as described above for HPS complementation assays with the following additional steps. After centrifugation at 16,000 x g for 10 min, supernatant was passed through a 0.2 μm filter to completely remove any residual cell or particulate material, and 0.5 mL of filtered medium was concentrated to ∼50 μl in a speedvac vacuum centrifuge. For SDS-PAGE analysis, these samples were separated on a 4-20% SDS-PAGE gel and subsequently detected by silver staining. For proteomic analysis, the concentrated medium was subsequently digested using the SP3 method (24). Briefly, 200 µg carboxylate-functionalized speedbeads (Cytiva Life Sciences) were added followed by the addition of acetonitrile to 80% (v/v) inducing binding to the beads. The beads were washed twice with 80% (v/v) ethanol and twice with 100% acetonitrile. Proteins were digested in 50 mM Tris-HCl, pH 8.5, with 0.5 µg Lys-C/Trypsin (Promega) and incubated at 37°C overnight. Tryptic peptides were desalted with the addition of 95% (v/v) acetonitrile binding the peptides back to the beads and washed once with 100% acetonitrile. Peptides were collected from the beads with two elutions of 1% (v/v) trifluoroacetic acid, 3% (v/v) acetonitrile. Cleaned-up peptides were then dried in a speedvac vacuum centrifuge and stored at −20°C until analysis.

For mass spectrometry analysis, tryptic peptides were suspended in 3% (v/v) ACN, 0.1% (v/v) trifluoroacetic acid (TFA) and directly injected onto a reversed-phase C18 1.7 µm, 130 Å, 75 mm X 250 mm M-class column (Waters), using an Ultimate 3000 nanoUPLC (Thermo Scientific). Peptides were eluted at 300 nL/minute with a gradient from 2% to 20% ACN in 40 minutes then to 40% ACN in 5 minutes and detected using a Q-Exactive HF-X mass spectrometer (Thermo Scientific). Precursor mass spectra (MS1) were acquired at a resolution of 120,000 from 350 to 1550 m/z with an automatic gain control (AGC) target of 3E6 and a maximum injection time of 50 milliseconds. Precursor peptide ion isolation width for MS2 fragment scans was 1.4 m/z, and the top 12 most intense ions were sequenced. All MS2 spectra were acquired at a resolution of 15,000 with higher energy collision dissociation (HCD) at 27% normalized collision energy. An AGC target of 1E5 and 100 milliseconds maximum injection time was used. Dynamic exclusion was set for 5 seconds. Rawfiles were searched against the Uniprot *Nostoc punctiforme* database UP000001191 downloaded 6/15/2023 using MaxQuant v.2.0.3.0. Cysteine carbamidomethylation was considered a fixed modification, while methionine oxidation and protein N-terminal acetylation were searched as variable modifications. All peptide and protein identifications were thresholded at a 1% false discovery rate (FDR). Statistical analysis was performed on cyclic loess normalized log2 transformed iBAQ and LFQ intensities using limma (25).

### Microscopy

Light and fluorescence microscopy was performed with an EVOS M5000 fluorescence microscope (Life Technologies) equipped with a 10x, 40x, or 63x objective lens. Excitation and emission were as follows: EVOS™ light cube, GFP (AMEP4651: excitation 470+/-22 nm, emission 525+/-50 nm) for UEA-fluorescein labeled HPS; EVOS™ Light Cube, DAPI (AMEP4650: excitation 357+/-44 nm, emission 447+/-60 nm) for immunofluorescence labeled PilA; and EVOS™ Light Cube, RFP (AMEP4652: excitation 531+/-40 nm, emission 593+/-40 nm) for cellular autofluorescence.

### Scytonemin production assay

The capacity of test strains to produce scytonemin was determined with a standard UV-A induction assay. Strains were grown in liquid 50% BG11_0_ medium under 12 h light (7 Wm^-2^), 12 h dark cycle and then placed under constant white light (13 Wm^-2^) amended with UVA radiation (3 W m^-2^) for seven days, after which biomass was collected by centrifugation, weighed, and then lipid soluble pigments extracted in 90% acetone for at least 24 h in the dark at 4°C, after which they were vortexed and centrifuged once more, discarding the pellet. Absorbance spectra of supernatants were recorded on a UV-visible spectrophotometer (Shimadzu UV-1601) between 350 and 800 nm, pigment concentrations resolved in the mixture using the equations of (26) developed for *Nosto*c extracts. Cell pigment contents were then expressed on a per wet weight basis. In all tests, the wild-type strain was assayed in parallel as a positive control.

The accumulation of scytonemin monomer in strains with disrupted scytonemin production was determined, as needed, through HPLC separation of extracts in a Waters e2695 equipped with a Supelco Discovery HS F5-5 column connected to a Waters 2998 PDA UV-Vis diode array detector using previously described protocols (27). Chlorophyll *a*, scytonemin and its monomer were monitored in the chromatograms at 407 nm, but spectra were recorded continuously between 200-800 nm using the PDA detector. Individual compounds were identified by their characteristic absorption maxima and appropriate retention time against true standards.

## Results

### The crtB locus

In *Nostoc punctiforme*, the *crtB* gene is encoded at a genetic locus (Fig. 1A) that contains genes implicated in HPS production via genomic co-occurrence and gene expression analysis (12). Three of these genes encode homologs of Wzx/Wzy-type polysaccharide synthesis and export system components (Wzy, Wza, and Wzc) (28). Immediately upstream and in the same orientation as *crtB* is a gene that codes for a putative aminotransferase (Npun_F0455, pfam01041), while immediately downstream, and in the same orientation is a gene that codes for a protein annotated as a CrtB-associated protein (Npun_F0457, TIGR04533). The function of this protein is unknown, but it is frequently found immediately downstream of *crtB* in the genomes of other cyanobacteria that possess a *crtB* gene. All of these genes are transcriptionally upregulated in developing hormogonia, with expression of *crtB* specifically dependent on the hormogonium sigma factor SigJ (29).

**Figure 1.**
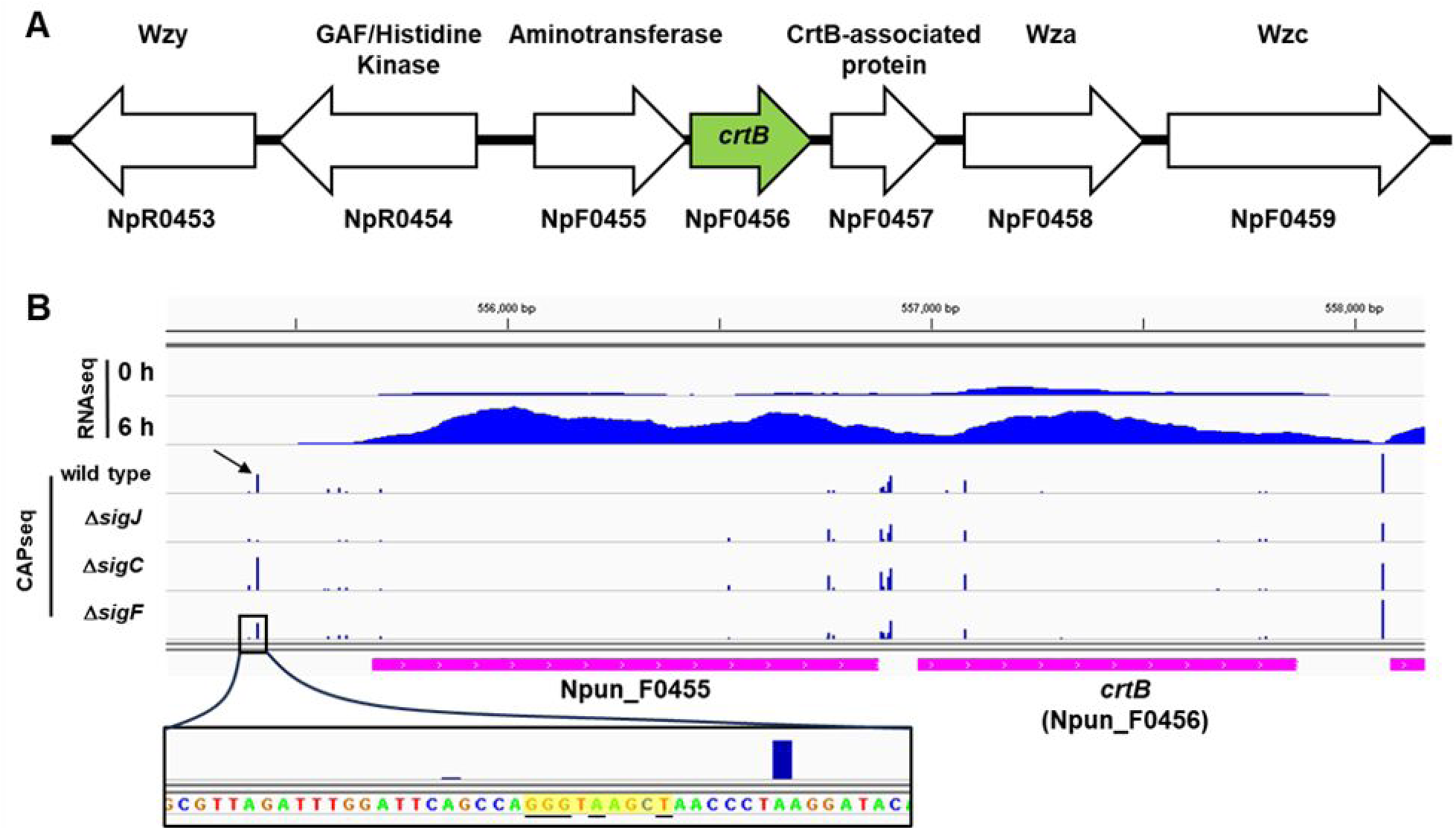
Architecture and gene expression of the *crtB* locus. (**A**) Schematic diagram depicting the *crtB* locus. Arrows represent protein coding genes, locus tags for each gene are listed underneath and putative proteins produced above. (B) Read map coverage from RNAseq and CAPseq data for various strains and time points as indicated. Read map coverage values for RNAseq and CAPseq indicated in brackets. Arrow indicates position of SigJ-dependent TSS. Inset below depicts sequence surrounding SigJ-dependent TSS, with J-Box highlighted in yellow, and absolutely conserved nucleotides of J-Boxes underlined.

To determine if *crtB* might be co-transcribed as part of an operon with the genes immediately upstream or downstream, we investigated previously published RNAseq (29) and Cappable-seq (CAPseq) (30) data sets to determine transcriptional start sites and total read coverage for this genomic region (Fig. 1B). CAPseq indicated the presence of a transcriptional start site (TSS) 272 bp upstream of the Npun_F0455 start codon that was previously identified as being SigJ-dependent (30). The −10 region for this TSS contains a consensus J-Box, providing further evidence that this is a SigJ-dependent promoter. Read coverage from RNAseq between Npun_F0455 and *crtB* is largely contiguous, supporting the notion that Npun_F0455 and *crtB* are co-transcribed as an operon from the SigJ-dependent promoter upstream of Npun_F0455. It should be noted that CAPseq data indicate the possible presence of several putative TSS in the intergenic region between Npun_F0455 and *crtB*, as well as internal to the *crtB* gene. However, their expression is not affected in any of the hormogonium-specific sigma factor mutants studied before. Thus, while it is possible that *crtB* is also transcribed monocistronically, it is unlikely that this is involved in upregulation during hormogonium development. In contrast, a single TSS is present 20 bp upstream of Npun_F0457 that was previously classified as SigC-dependent (30) and read coverage from RNAseq tapers off substantially in the intergenic region between *crtB* and Npun_F0457. These results indicate that Npun_F0457 is unlikely to be transcribed as part of an operon with *crtB*.

### CrtB is essential for motility

The increased expression of *crtB* in developing hormogonia indicates that it may play a key role in hormogonia development and motility. To test this hypothesis, a strain with an in-frame deletion of *crtB* was constructed. Deletion of *crtB* resulted in the loss of motility as gauged both by plate motility assays of colony spreading and time-lapse microscopy of individual filaments following hormogonium induction (Fig. 2A, SMOV 1). Despite the absence of motility, the Δ*crtB* strain produced morphologically distinct hormogonia, indicating that *crtB* does not influence early stages of hormogonium development. Immunoblot analysis showed that, compared to the wild-type strain, the Δ*crtB* strain produces equivalent amounts of both the major Pilin PilA and the methy-accepting chemotaxis protein HmpD.

**Figure 2.**
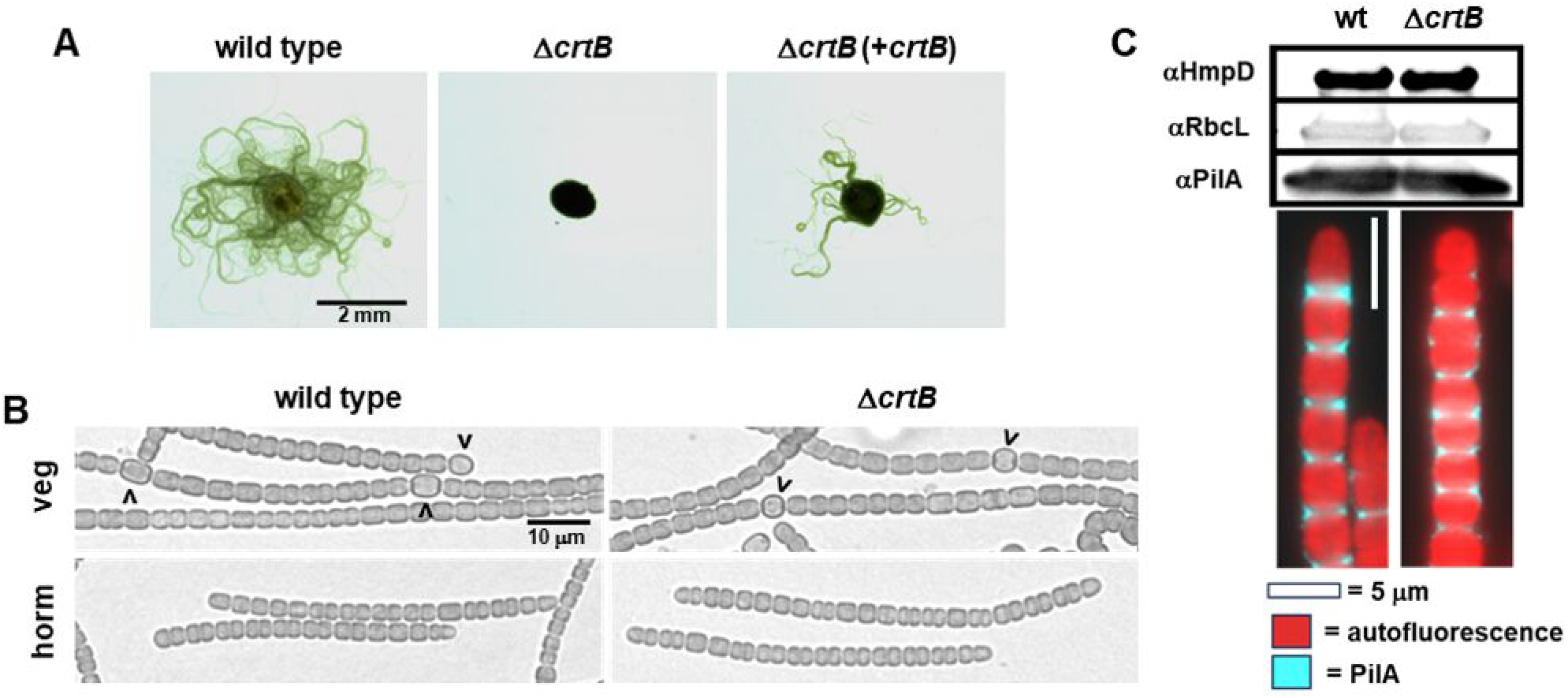
Hormogonium development and motility of the Δ*crtB* strain. (A) Plate motility assays of the wild type, Δ*crtB*, and complemented Δ*crtB* strain (+*crtB*) as indicated. (B) Light micrographs of vegetative (veg) and hormogonium (horm) filaments 18 h following hormogonium induction. Carets indicate heterocysts. (C) Immunoblot and immunofluorescence analysis of the hormogonium-specific proteins HmpD and PilA for strains as indicated. RbcL serves as a protein loading control.

Because expression of *pilA* and *hmpD* is regulated by distinct branches of the hormogonium gene regulatory network (29), this suggests that *crtB* does not have a substantial influence on the hormogonium developmental program. Furthermore, immunofluorescence analysis demonstrated that PilA was present on the surface of the Δ*crtB* strain, suggesting that the T4P systems retain function in that it can export PilA out of the cell.

### CrtB influences HPS accumulation and biofilm formation

Considering that *crtB* is found at a locus with other genes implicated in the production of HPS, it is a possibility that *crtB* also plays a role in this process. This would be consistent with the loss of motility in the Δ*crtB* strain shown above. To test this hypothesis, we studied HPS production through lectin-blot analyses (Fig. 3A). This showed that the amounts of soluble HPS released into the medium by wild type and Δ*crtB* strains was comparable although more variable between replicates for Δ*crtB*. In contrast, cell-associated HPS content, analyzed by fluorescent lectin staining, was substantially less in the Δ*crtB* strain than in the wild type. Moreover, the fluorescently labeled HPS that directly accumulated adjacent to filaments was patently reduced in the Δ*crtB* strain compared to that in the wild type (Fig. 3B). These results imply that while the Δ*crtB* strain can produce HPS, its ability to retain it in close proximity to the filaments is disrupted.

**Figure 3.**
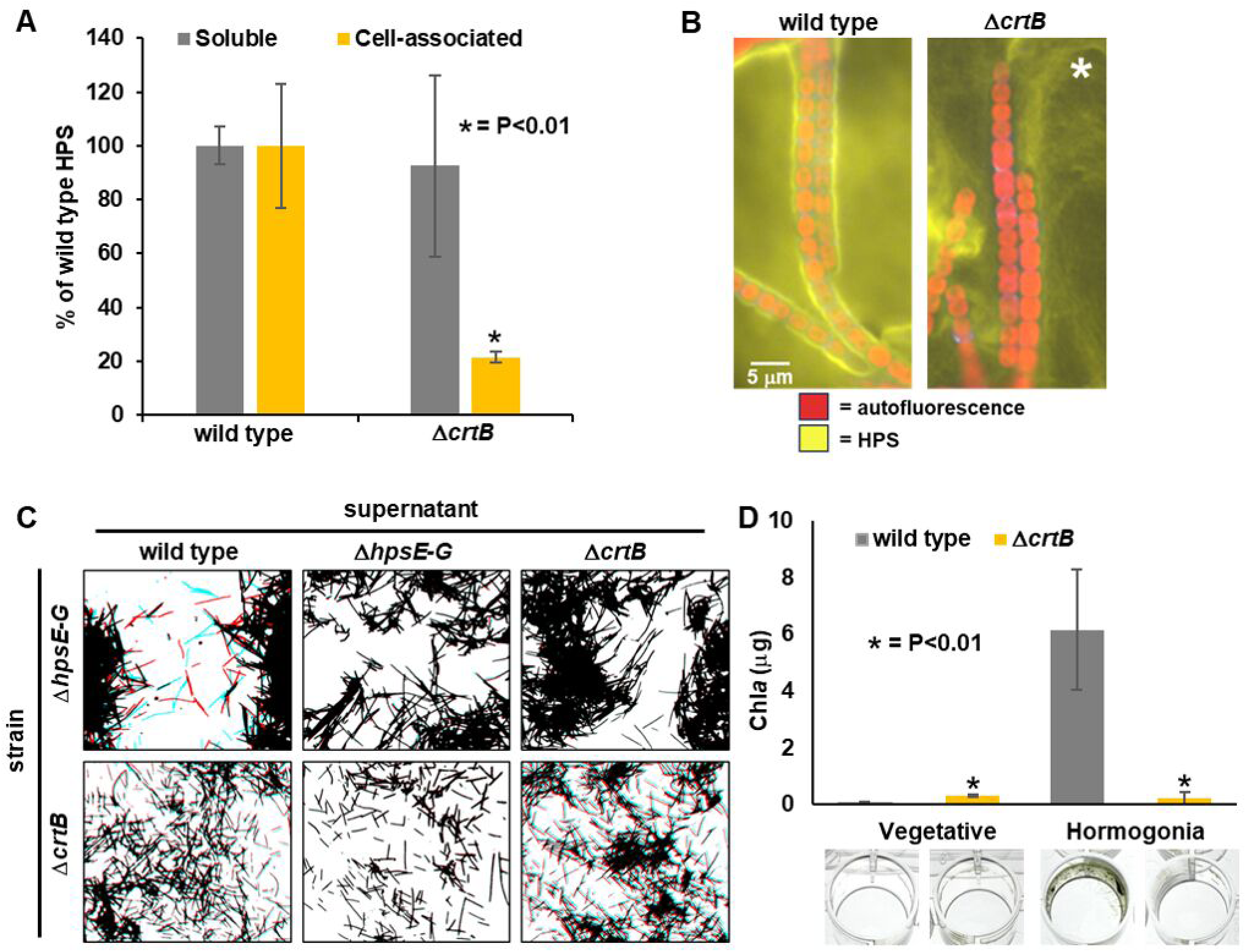
Production of HPS and biofilm formation in the Δ*crtB* strain. (**A**) Quantification of soluble and cell associated HPS by lectin-based analysis. Bars = +/- 1 S.D., p-value from student t-test between wild type and Δ*crtB* strains indicated for significant differences. (**B**) Fluorescence micrographs of lectin-stained HPS (strains as indicated). * Indicates that the brightness and contrast of the Δ*crtB* image were increased compared to the wild-type image to visualize HPS, which was much lower for cell-associated HPS in Δ*crtB*. (**C**) Exogenous complementation of mutant strains by addition of culture medium containing HPS. Various strains (as indicated) were incubated with cell-free culture supernatants from strains as indicated and subjected to time-lapse microscopy. Images depicted are derived from SMOV 2 and are a merge of the first frame of the time-lapse colored red, and the last frame colored blue. Non-motile filaments appear black due to overlap of red and blue channels, while motile filaments appear separately as blue and red. (**D**) Quantitative Biofilm assays (top) and representative images (bottom) of cultures containing vegetative or hormogonium filaments (strains as indicated). p-value derived from students t-test comparing either vegetative filaments or hormogonia between the wild type and Δ*crtB* strains.

For some previously characterized mutants that lose the ability to produce HPS, motility can be partially restored by exogenous addition of HPS (10, 12), while for other mutants, this is not the case (12). For the latter, the mutants typically also show defects in accumulation of surface PilA, indicating these genes alter both HPS production and T4P activity, which could explain the failure of exogenous HPS to restore motility (12). Given that the Δ*crtB* strain retains surface PilA and still produces abundant soluble HPS, the loss of motility could be explained if either the composition of the HPS had been altered in the Δ*crtB* strain so that it no longer adheres to the filament surface, or if the composition of the cell surface has been altered so that it no longer adheres to the HPS. To test these alternative hypotheses, the ability of HPS produced by the Δ*crtB* strain to restore motility in an HPS-deficient mutant was tested, as was the ability of wild type HPS to restore motility to the Δ*crtB* mutant (Fig. 3C, SMOV. 2). As previously reported, the addition of supernatant from medium containing wild-type hormogonia restored motility in the HPS-deficient Δ*hpsE-G* strain (10). In contrast, supernatant from the Δ*crtB* strain failed to restore motility to the Δ*hpsE-G* strain, despite containing equivalent levels of HPS based on lectin-blotting. This result is consistent with the hypothesis that the composition of HPS may be altered in the Δ*crtB* strain in such a way that it fails to support motility. Conversely, supernatant from medium containing wild-type hormogonia also failed to restore motility in the Δ*crtB* strain, consistent with the hypothesis that this strain may have an alteration in the filament surface resulting in defective adhesion to HPS. These results indicate that *crtB* may affect both the composition of the HPS and the cell surface to facilitate adhesion of HPS and motility.

We have routinely observed that mutants of *N. punctiforme* that lose motility also fail to aggregate (3) and form biofilms on the surface of culture vessels in liquid cultures (6), although we have not documented the latter for many of the characterized motility deficient mutants. Moreover, biofilm formation has also been shown to be positively correlated with motility in the filamentous cyanobacterium *Leptolyngbya boryana* (5). To determine what role hormogonia and motility, and *crtB*, may play in biofilm formation, the ability of both wild-type and Δ*crtB* vegetative and hormogonium filaments to form biofilms was determined (Fig. 3D). Vegetative filaments of both the wild type and Δ*crtB* strain failed to display obvious biofilm formation on the side walls or bottoms of 24 well plates, although the Δ*crtB* mutant did tend to accumulate a ring of filaments that adhered to the side wall at the air-liquid interface to a slightly greater extent than the wild type. Wild-type hormogonium filaments accumulated biofilms that were apparent primarily on the side walls of these plates well below the air-liquid interface. This was not the case for hormogonium filaments of the Δ*crtB* strain. Thus, in additionto its critical role in motility, *crtB* plays an important role in the formation of biofilms under these conditions.

### CrtB affects the presence of PEP-CTERM proteins in the exoproteome

CrtB is predicted to process secreted proteins containing a PEP-CTERM domain, facilitating their anchoring to the outer membrane or their release from the cell. Based on this notion, the deficiencies in motility and biofilm formation observed for the Δ*crtB* mutant are most likely due to a failure to properly process and sort these PEP-CTERM proteins. The genome of *N. punctiforme* harbors 35 genes coding for proteins containing putative PEP-CTERM domains (Fig. 4A). Seven of these genes are substantially upregulated in developing hormogonia (Fig. 4A) (29). Five of these, Npun_F0296, Npun_F4801, Npun_ER021, Npun_R0434, and Npun_R3960, show a remarkably similar expression pattern in developing wild-type hormogonia, as well as in hormogonium-specific sigma factor deletion strains, indicating they are primarily dependent on SigJ for enhanced transcription in hormogonia. However, previous work has indicated that these genes lack a consensus J-Box in the −10 region of their promoters (29, 30), implying that regulation by SigJ is indirect. In contrast to these five PEP-CTERM genes, the other two genes upregulated in hormogonia, Npun_R4127 and Npun_R6196, display different expression profiles. Given their enhanced expression in developing hormogonia, it is possible that these seven PEP-CTERM proteins play a critical role in motility and biofilm formation.

**Figure 4.**
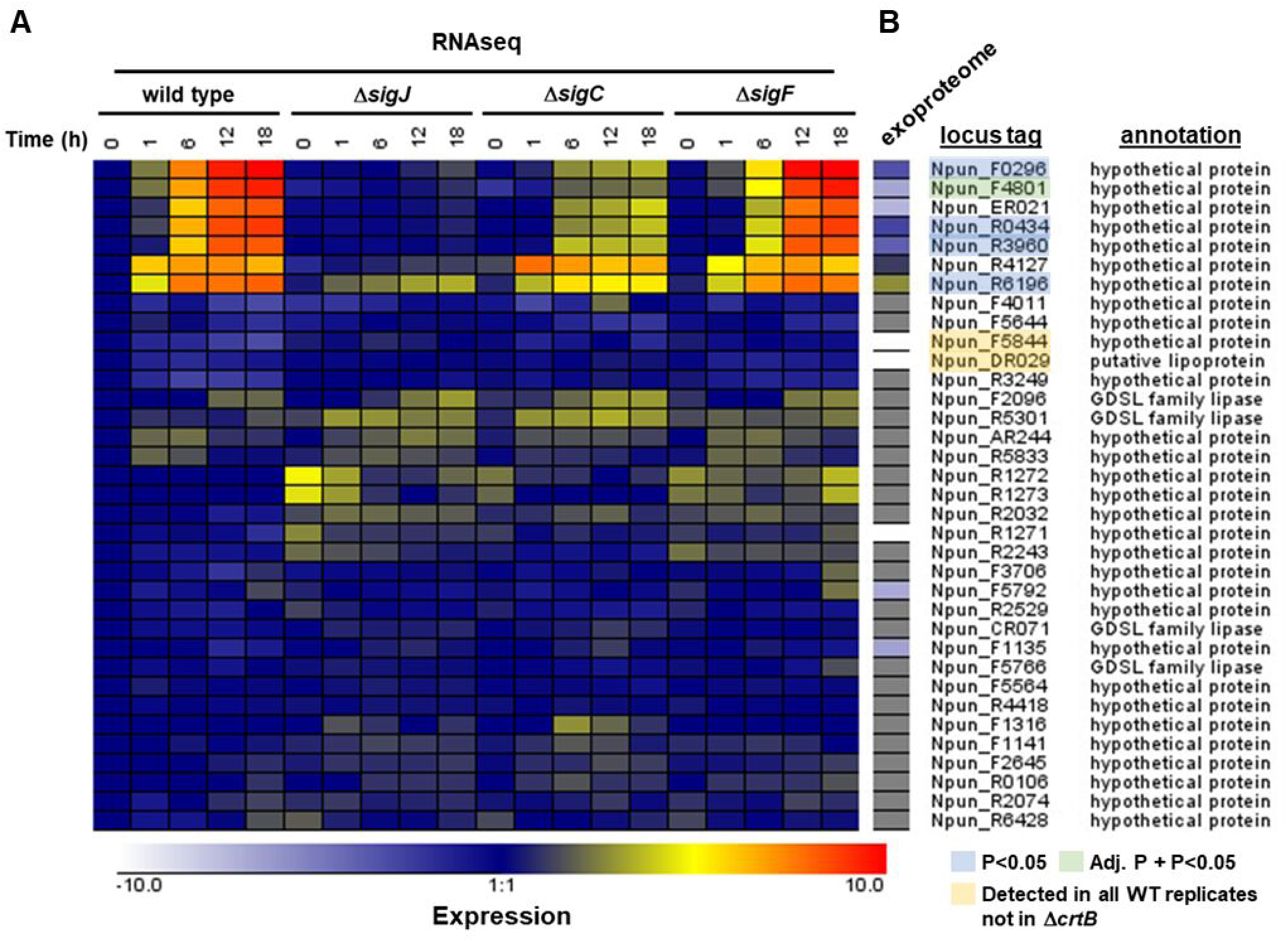
Gene expression and exoproteome analysis of putative PEP-CTERM proteins in developing hormogonia. (**A**) RNAseq based transcription profiles of putative PEP-CTERM proteins of the wild type, and sigma factor mutant strains (as indicated). Expression = Log2(Exp/WT t=0). (**B**) Mass-spectrometery based quantitative analysis of PEP-CTERM protein abundance in the wild type and Δ*crtB* strain. Expression = Log2(Δ*crtB* /WT). Grey indicates that the proteins were not detected.

To test this hypothesis, the exoproteome of hormogonia was analyzed. SDS-PAGE followed by silver staining of culture medium for both the wild type and Δ*crtB* strain indicated that the exoproteome for each was largely similar, although there were at least two distinct proteins present in the wild type that appeared to be absent or substantially diminished in the Δ*crtB* mutant (Fig. S1). Proteomic analysis of these samples by mass-spectrometry detected 12 of the 35 PEP-CTERM proteins encoded in the *N. punctiforme* genome in the exoproteome of the wild-type strain, including all 7 of those specifically upregulated in developing hormogonia (Fig. 4B, Data Set S1). A single protein was considered differentially abundant in the Δ*crtB* strain vs the wild type by both p-value (7.98e-^06^) and adjusted p-value (0.0105), the PEP-CTERM protein Npun_F4801, which was reduced ∼92 fold in the Δ*crtB* strain. This protein is encoded by one of the five PEP-CTERM genes that exhibit a similar indirect SigJ-dependent transcriptional profile in developing hormogonia. Levels of the other proteins encoded by this gene set were also substantially diminished in the Δ*crtB* mutant, with three of the proteins considered significantly diminished by p-value (p<0.05) but not adjusted p-value (Fig. 4B, Data Set S1). The low statistical confidence is likely a result of substantial variation that was observed between replicates. In contrast, for the other two PEP-CTERM proteins encoded by genes upregulated in hormogonia, Npun_R4127 and Npun_R6196, protein abundance was increased in the Δ*crtB* strain ∼2.3 (p-value 0.151) and ∼6.7 (p-value 0.025) fold respectively.

Notably, none of the predicted molecular weights for the hormogonium-expressed PEP-CTERM proteins corresponds to the apparent molecular weight of the two proteins missing or reduced in the Δ*crtB* exoproteome in the SDS-PAGE analysis, although it is conceivable that this could be due to substantial post-translational modification of these PEP-CTERM proteins, in particularly glycosylation. The other 5 PEP-CTERM proteins detected in the exoproteome of the wild type were all reduced in the Δ*crtB* strain, with two detected in all three replicates of the wild type and none of the three replicates for the Δ*crtB* strain (Fig. 4B). Given that CrtB would be expected to remove the PEP-CTERM domain from processed proteins, we also analyzed the mass-spectrometry data to determine if there was differential detection of peptides covering the PEP-CTERM domain between the wild type and Δ*crtB* mutant. However, peptide coverage of the PEP-CTERM domain for these proteins was never detected in either strain (Fig. S2). Collectively, these results are consistent with the hypothesis that CrtB is involved in the sorting of PEP-CTERM proteins, and that the defects in motility and biofilm formation for the Δ*crtB* strain are at least partly due to alteration of extracellular PEP-CTERM proteins.

### CrtB is essential for production of scytonemin

Many cyanobacteria, including *N. punctiforme*, can produce the UV-absorbing pigment scytonemin to protect cells from UVA exposure. In *N. punctiforme* two genetic loci are known to be involved in scytonemin production, the *scy locus,* largely biosynthetic, and the *ebo* locus, involved with transporting scytonemin monomers to the periplasm, where late biosynthetic steps occur (19, 27, 31). Three genes within the *scy* locus encode putative PEP-CTERM proteins, *scyD* (Npun_R1273) *scyE* (Npun_R1272) and *ScyF* (Npun_R1271). Deletion of *scyE* abolished scytonemin production, leading to the accumulation of a scytonemin intermediate in the periplasm, while a *scyF*-deletion strain exhibited reduced scytonemin accumulation (19). The deletion of *scyD* had no apparent phenotypic effect on scytonemin biosynthesis (19). The critical role of putative PEP-CTERM proteins in scytonemin production implies that CrtB may also be essential for this process, presuming that processing by CrtB is essential for Scy PEP-CTERM protein function. Notably, ScyF was detected in two of three replicates of the hormogonium exoproteome of the wild-type strain, but none of the three replicates for Δ*crtB*, even when expression of the *scy* locus is suppressed during hormogonia formation and uninduced without exposure to UVA (32). This provides direct experimental evidence that CrtB may be involved in processing the Scy PEP-CTERM proteins and could indicate that CrtB is also essential for scytonemin production.

To test this hypothesis, the Δ*crtB* mutant was exposed to UVA radiation and scytonemin production quantified (Fig. 5A). In the wild-type strain, as expected in this UVA inducible system, scytonemin accumulated to ∼0.6 mg/g cell material (wet weight) under UVA exposure and was undetectable in the absence of UVA. In contrast, the Δ*crtB* strain failed to exhibit detectable scytonemin in either the presence or absence of UVA. As with motility, scytonemin production could be complemented by the reintroduction of *crtB* in trans. The Δ*crtB* strain also exhibited a reduction in the amount of Chl *a* compared to the wild type, and this was more pronounced under UVA exposure (Fig. S3). This data indicates that CrtB plays an essential role in the production of scytonemin. We also compared Δ*crtB* to Δ*scyE,* a previously described scytoneminless mutant that accumulates the scytonemin monomer in the periplasm during UVA induction, as the monomer condensation reaction is disabled. Unlike Δ*scyE,* and unlike mutants in the *ebo* locus, Δ*crtB* did not seem to accumulate the monomer to clearly detectable levels (Fig 5B).

**Figure 5.**
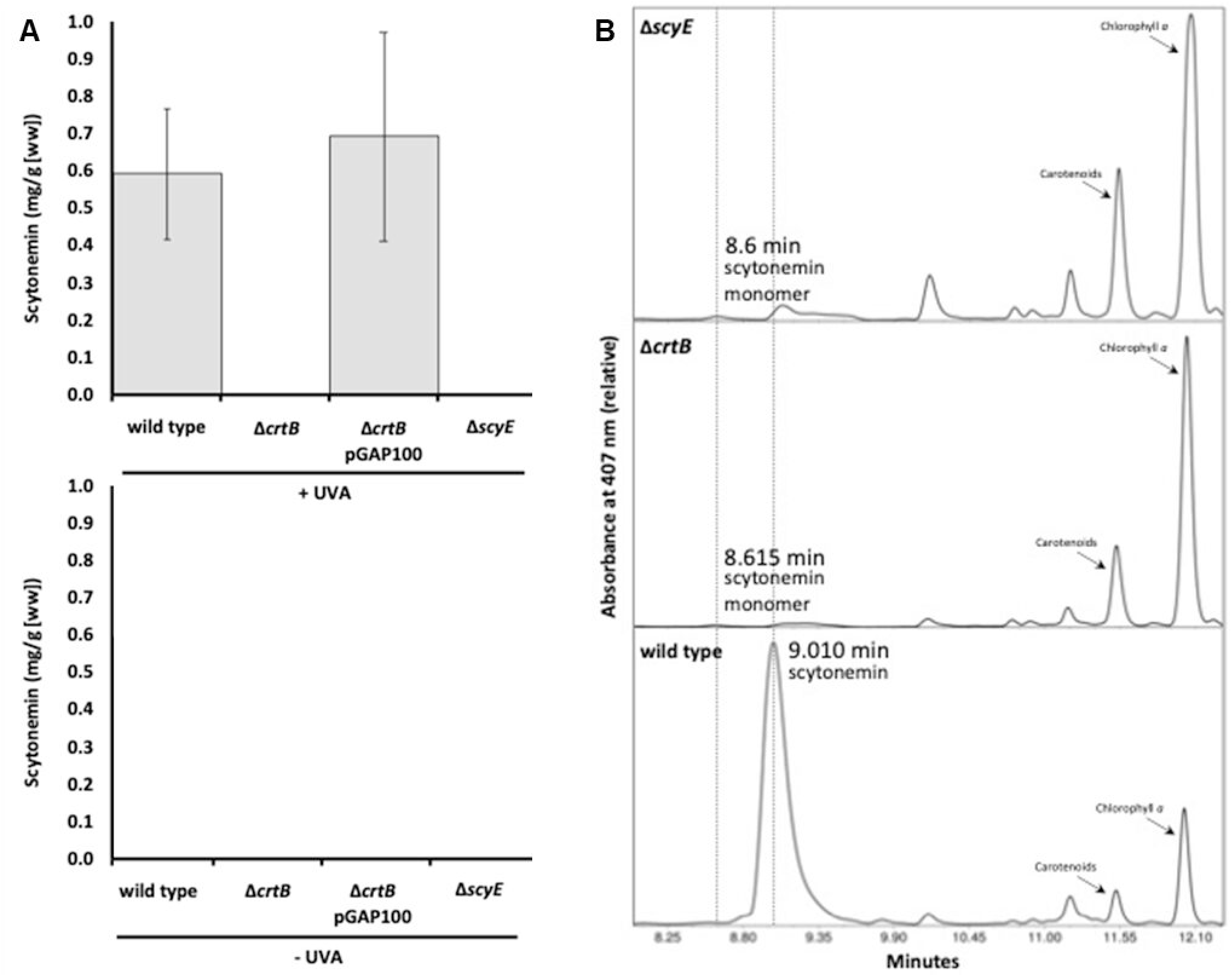
Effect of CrtB on scytonemin production. (A) Scytonemin production in strains (as indicated) in the presence and absence of UVA exposure. Error bars = +/-1 S.D. (B) Separation and characterization of scytonemin and the monomer precursor accumulated after UVA induction in various strains (as indicated) by HPLC, showing production of scytonemin monomer at 8.615 min and scytonemin at 9.010 min.

## Discussion

Figure 6 depicts a working model for the role of the cyanoexosortase B systems in motility and scytonemin production in *N. punctiforme*. In the case of motility (Fig. 6A), a subset of PEP-CTERM proteins is specifically expressed in developing hormogonia. These proteins are transported across the cytoplasmic membrane via the general secretory pathway and the N-terminal signal peptide is removed. The PEP-CTERM domain tethers the protein to the cytoplasmic membrane until it is processed by CrtB, removing the PEP-CTERM domain, and possibly covalently linking the protein to HPS or some other molecule that is transported across the outer membrane. How the proteins transverse the outer membrane is currently unclear, but one possibility is that they transit along with the HPS via a dedicated polysaccharide secretion system. Notably, the *crtB* locus encodes a Wza-type outer membrane auxiliary (OMA) family protein (Npun_F0459), and these are known to be involved in translocation of polysaccharide across the outer membrane. It is also possible that the CrtB-associated protein encoded immediately downstream of *crtB* (Npun_F0457) may play a role in this process. However, more work is required to verify this hypothesis and definitively determine the route of exit for PEP-CTERM proteins across the outer membrane. Once outside the cell these proteins facilitate interaction between the cell surface and HPS promoting tight adhesion and thereby facilitating locomotion. It is possible that the PEP-CTERM proteins may be covalently anchored to the outer membrane, the HPS, or both. The failure of wild-type HPS to restore motility to the Δ*crtB* mutant implies that there is a cell-surface defect in the Δ*crtB* mutant reducing adhesion to HPS and indicating that PEP-CTERM proteins are attached to the outer membrane. Conversely, the failure of HPS from the Δ*crtB* strain to restore motility in the Δ*hpsE-G* mutant, despite apparently similar levels of HPS to the wild type, indicates an alteration in the composition of the HPS from the Δ*crtB* strain, and supports the idea that PEP-CTERM proteins are a component of HPS. This is consistent with the theory that PEP-CTERM proteins are anchored to both the outer membrane and the HPS, and that this is essential for motility, but more work will be needed to validate this hypothesis. The loss of adhesion between extracellular polysaccharides and the cell surface could also explain the loss of biofilm formation in the Δ*crtB* strain. This model is also consistent with work in other bacteria indicating that exosortases and PEP-CTERM proteins are critical for stabilizing the interaction between the cell surface and EPS (16, 17).

**Figure 6.**
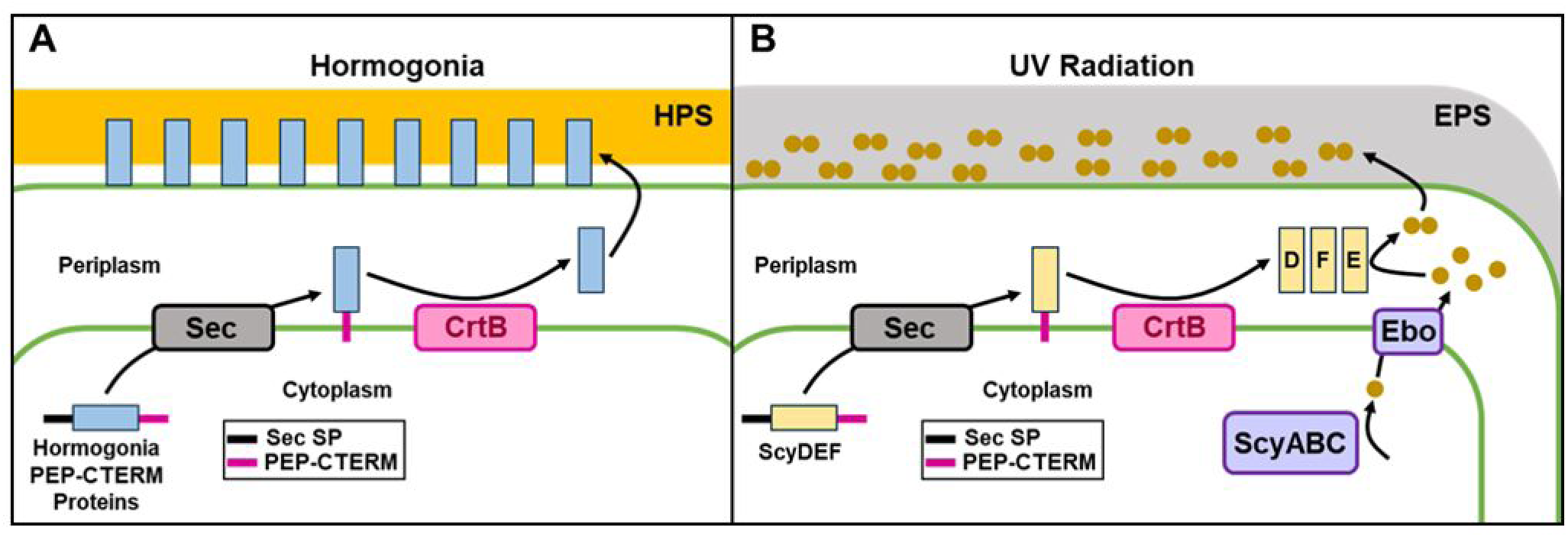
A working model depicting the role of CrtB and PEP-CTERM proteins in motility and scytonemin biosynthesis. (A) Hormogonium PEP-CTERM proteins are transported across the cytoplasmic membrane via the general secretory (Sec) pathway and the N-terminal sec signal peptide (Sec-SP) is removed, after which the C-terminal PEP-CTERM domain anchors the proteins to the cytoplasmic membrane until it is processed by CrtB. The mature PEP-CTERM proteins are then transported across the outer membrane via an undefined pathway where they facilitate adhesion between the HPS and the cell surface. (B) The scytonemin-specific PEP-CTERM proteins ScyD, ScyE, and ScyF are transported across the cytoplasmic membrane and processed as described in A. Scytonemin monomers are synthesized in the cytoplasm by ScyABC, the monomers are transported across the cytoplasmic membrane by the Ebo system, and mature ScyE dimerizes the monomers to produce scytonemin, which is subsequently transported across the outer membrane via an undefined pathway where it associates with extracellular polysaccharides (EPS).

For scytonemin production, the ScyD, ScyE, and ScyF proteins are secreted across the cytoplasmic membrane via the sec pathway and subsequently processed by CrtB in a similar manner to the hormogonium PEP-CTERM proteins. At this point it is not clear whether ScyD, ScyE, and ScyF are attached to another molecule and/or exported across the outer membrane or whether they are just released into the periplasm. Previous work has verified that ScyE and ScyF are abundant in the periplasm (27), but there is no data currently to support association of these proteins with the outer membrane. Scytonemin monomers are synthesized in the cytoplasm by ScyABC, transported across the cytoplasmic membrane by the Ebo system, and the final enzymatic step to dimerize the monomers and produce scytonemin is performed by ScyE either in the periplasm or on the cell surface. It is clear from our results that CrtB is necessary for scytonemin synthesis, and we would have expected the scytonemin monomers to have accumulated under UVA radiation, as the Ebo system for monomer transport was intact. That this prediction did not materialize may simply be the result of a secondary effect of a leaky outer membrane, consistent with significant pleiotropic effects on cell morphology. However, more direct effects of CrtB involving the necessity of protein excretion to organize the Ebo systems cannot be discarded at this point. While the final destination for the Scy PEP-CTERM proteins is unresolved, scytonemin is clearly either transported across the outer membrane following dimerization, or monomers transverse the outer membrane and are subsequently dimerized on the exterior of the cell, as mature scytonemin is found outside of the cell, associated with extracellular polysaccharides (33). The association of scytonemin with EPS is also notable, as it fits a consistent theme of exosortases and PEP-CTERM proteins being related to processes associated with accumulation of EPS. The necessity of CrtB in scytonemin production also has important implications for efforts to genetically engineer non-native hosts to produce scytonemin. Currently, such efforts have only led to the production of scytonemin monomers (34), and the data presented here imply that introduction of *crtB* into non-native hosts may be essential for production of mature scytonemin.

There are a total of 35 predicted PEP-CTERM proteins encoded in the *N. punctiforme* genome. While the results reported here provide a plausible explanation for the role of a subset of these in motility and scytonemin production, the function of the other PEP-CTERM proteins remains cryptic and suggests that the CrtB system may be involved in other biological functions. The hormogonium exoproteome analysis detected several additional PEP-CTERM proteins, most notably Npun_F5844 and Npun_DR029, which were detected in all three replicates for the wild type but none for Δ*crtB*, although at low peptide coverage (1 and 2 respectively). This indicates that at least some of these other PEP-CTERM proteins are expressed and processed by CrtB. Expression for both genes encoding these proteins is also downregulated in developing hormogonia implying that they perform a function in vegetative cells rather than hormogonia. *N. punctiforme* is known to produce a variety of EPS aside from the HPS associated with hormogonia and motility, such as the mannose-rich EPS produces in the aseriate growth state observed in cultures containing sucrose and in symbiotic association with the hornwort *Anthoceros punctatus* (21). Given the evidence presented here that CrtB and PEP-CTERM proteins are involved in the EPS related processes of motility, biofilm formation, and scytonemin production, as well as the association of exosortases and PEP-CTERM proteins with EPS related functions in other bacteria (13, 14), investigating the role of the CrtB system in other EPS-related phenotypes might be a useful starting point for defining what, if any, other roles this system plays in cyanobacteria.

## Acknowledgements

The authors would like to acknowledge the following funding sources: the work of C.C.E. was supported by NIH S10-OD025267; the work of D.H.H. was supported by the National Center for Biotechnology Information of the National Library of Medicine (NLM), National Institutes of Health; the work of D.D.R. was supported by NSF-IOS 2420339.

## Conflict of Interest

The authors declare that there are no conflicts of interest.

